# Enhanced food motivation in obese mice is controlled by D1R expressing spiny projection neurons in the nucleus accumbens

**DOI:** 10.1101/2022.01.12.476057

**Authors:** Bridget A Matikainen-Ankney, Alex A Legaria, Yvan M Vachez, Caitlin A Murphy, Yiyan Pan, Robert F Schaefer, Quinlan J McGrath, Justin G Wang, Maya N Bluitt, Aaron J Norris, Meaghan C Creed, Alexxai V Kravitz

## Abstract

Obesity is a chronic relapsing disorder that is caused by an excess of caloric intake relative to energy expenditure. In addition to homeostatic feeding mechanisms, there is growing recognition of the involvement of food reward and motivation in the development of obesity. However, it remains unclear how brain circuits that control food reward and motivation are altered in obese animals. Here, we tested the hypothesis that signaling through pro-motivational circuits in the core of the nucleus accumbens (NAc) is enhanced in the obese state, leading to invigoration of food seeking. Using a novel behavioral assay that quantifies physical work during food seeking, we confirmed that obese mice work harder than lean mice to obtain food, consistent with an increase in the relative reinforcing value of food in the obese state. To explain this behavioral finding, we recorded neural activity in the NAc core with both *in vivo* electrophysiology and cell-type specific calcium fiber photometry. Here we observed greater activation of D1-receptor expressing NAc spiny projection neurons (NAc D1^SPNs^) during food seeking in obese mice relative to lean mice. With *ex vivo* slice physiology we identified both pre- and post-synaptic mechanisms that contribute to this enhancement in NAc D1^SPN^ activity in obese mice. Finally, blocking synaptic transmission from D1^SPNs^ decreased physical work during food seeking and attenuated high-fat diet-induced weight gain. These experiments demonstrate that obesity is associated with a selective increase in the activity of D1^SPNs^ during food seeking, which enhances the vigor of food seeking. This work also establishes the necessity of D1^SPNs^ in the development of diet-induced obesity, establishing these neurons as a potential therapeutic target for preventing obesity.

## Introduction

Obesity is caused by an imbalance in caloric intake relative to expenditure. Recent work has appreciated the critical role of food reinforcement and food reward in obesity^1^. People with obesity will work harder to obtain food^2^, and individual levels of food reinforcement predict future weight gain^3^. The modern food environment affords near-constant access to calorie dense foods, so heightened food reinforcement can drive overeating and cause weight gain. Therefore, we set out to understand adaptations in neural circuits mediating food reinforcement in obese animals, and how these adaptations contribute to the vigor of food seeking and the propensity of individuals to gain weight.

The nucleus accumbens (NAc) is a critical mediator of food reinforcement and decision making in obesity^4,5^. Specifically, human neuroimaging studies have identified changes in the NAc and its afferents in people with obesity, leading to the enhanced impact of food-related information on these circuits^6–13^. Pre-clinical rodent research has further identified changes in synaptic plasticity and intrinsic excitability in the NAc of obese rodents^14–16^. Despite these links, the NAc cell types that are disrupted in obese animals remain unknown. The NAc comprises two populations of projection neurons, which express either the dopamine D1 or D2 receptor (termed: D1 or D2 receptor-expressing Spiny Projection Neurons, D1^SPNs^ and D2^SPNs^) ^17,18^. D1^SPNs^ invigorate and reinforce actions, while D2^SPNs^ generally inhibit them^19,20^, with the balance of activity between these pathways determining whether an animal engages in a behavior or not^20–22^. Based on their respective roles, we hypothesized that the balance of accumbal D1^SPN^:D2^SPN^ activity would be shifted towards elevated D1^SPN^ activity during food seeking in obese mice, and that this imbalance in D1^SPN^:D2^SPN^ activity would be necessary for the development of diet-induced obesity.

We tested our hypothesis with a combination of *in vivo* and *ex vivo* approaches. Supporting our hypothesis, we found that diet-induced obesity was associated with a selective enhancement of *in vivo* calcium activity of NAc D1^SPNs^ (and not D2^SPNs^) during food seeking, thereby shifting the balance of accumbal activity towards D1^SPNs^. These changes in D1^SPN^ activity were associated with increases in both the intrinsic excitability of D1^SPNs^ and excitatory synaptic drive onto D1^SPNs^ (and again not D2^SPNs^) as measured *ex vivo*. Finally, blocking synaptic output of D1^SPNs^ via viral expression of a tetanus toxin light-chain subunit reduced food seeking and attenuated high-fat diet induced weight gain, confirming the necessity of NAc D1^SPNs^ in the development of diet-induced obesity. Our data establish that obesity is associated with excitatory adaptations in NAc D1^SPNs^ that invigorate food seeking. We further conclude that NAc D1^SPNs^ mediate the development of diet-induced obesity, suggesting that they may represent an underexplored therapeutic target for treating obesity.

## Results

### Diet-induced obesity increases physical work during food seeking

It has been challenging to determine whether obese animals are more or less motivated to obtain food than lean animals: on one hand, obese rodents eat more than lean rodents when food is provided freely^23–25^; but on the other hand, they expend less effort to obtain food when they have to work for it^26–28^. This decrease in effort has primarily been observed in progressive ratio tasks that require increasing numbers of operant responses to obtain food, which we also observed (Suppl Fig 1). Poor performance of obese mice on the progressive ratio task has been interpreted as a decrease in their food motivation^27,29^. However, the progressive ratio task does not solely measure effort. Rather, this task conflates the *amount* of work performed (the number of responses) with the *time* it takes to complete this work (it takes longer to complete larger numbers of responses). This is an important point to consider, as delayed rewards are devalued more strongly by people with obesity^30,31^, so poor performance on progressive ratio tasks may reflect a decrease in food motivation *or* an unwillingness to work for delayed food rewards.

**Supplemental Figure 1.**
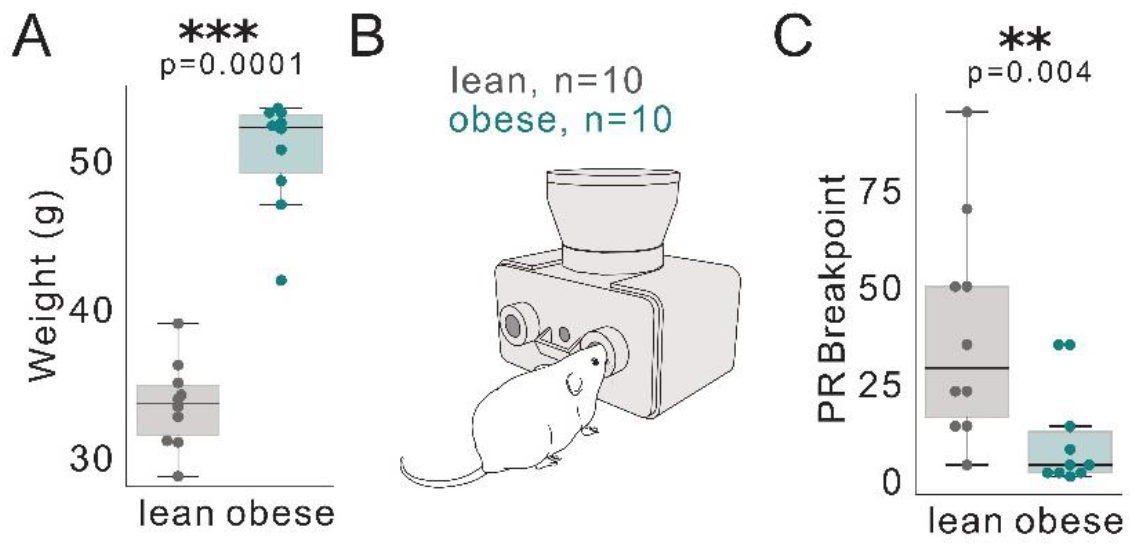
Obesity decreases frequency of responding for food rewards, (a) Obese mice weigh significantly more than lean. p=0.0001. (b) Obese and lean mice were trained to nosepoke on an active port to earn a food pellet, (c) Obese mice reach signficantly lower breakpoints in a progressive ratio task, p=0.004. Mann Whitney U tests to comapre means in a, c.

To address the confound caused by temporal delay, we developed a behavioral task to measure effort output and physical work during food seeking without the temporal confound of delaying the reward due to increased lever press frequency. In the designed task, mice pressed a non-moveable load-cell lever to obtain food pellets (Supplemental Video 1). The first pellet required 1g of force and each earned pellet increased this force requirement by 1g. Pellets were delivered immediately after completing the force requirement, thereby increasing the work requirement without introducing temporal delays. The force requirement reset to 1g when the mouse failed to earn a pellet in 30 minutes (Fig 1A), which allowed us to run the task in a “closed economy” design over multiple days, with animals earning all food from the task (Fig 1C).

**Figure 1.**
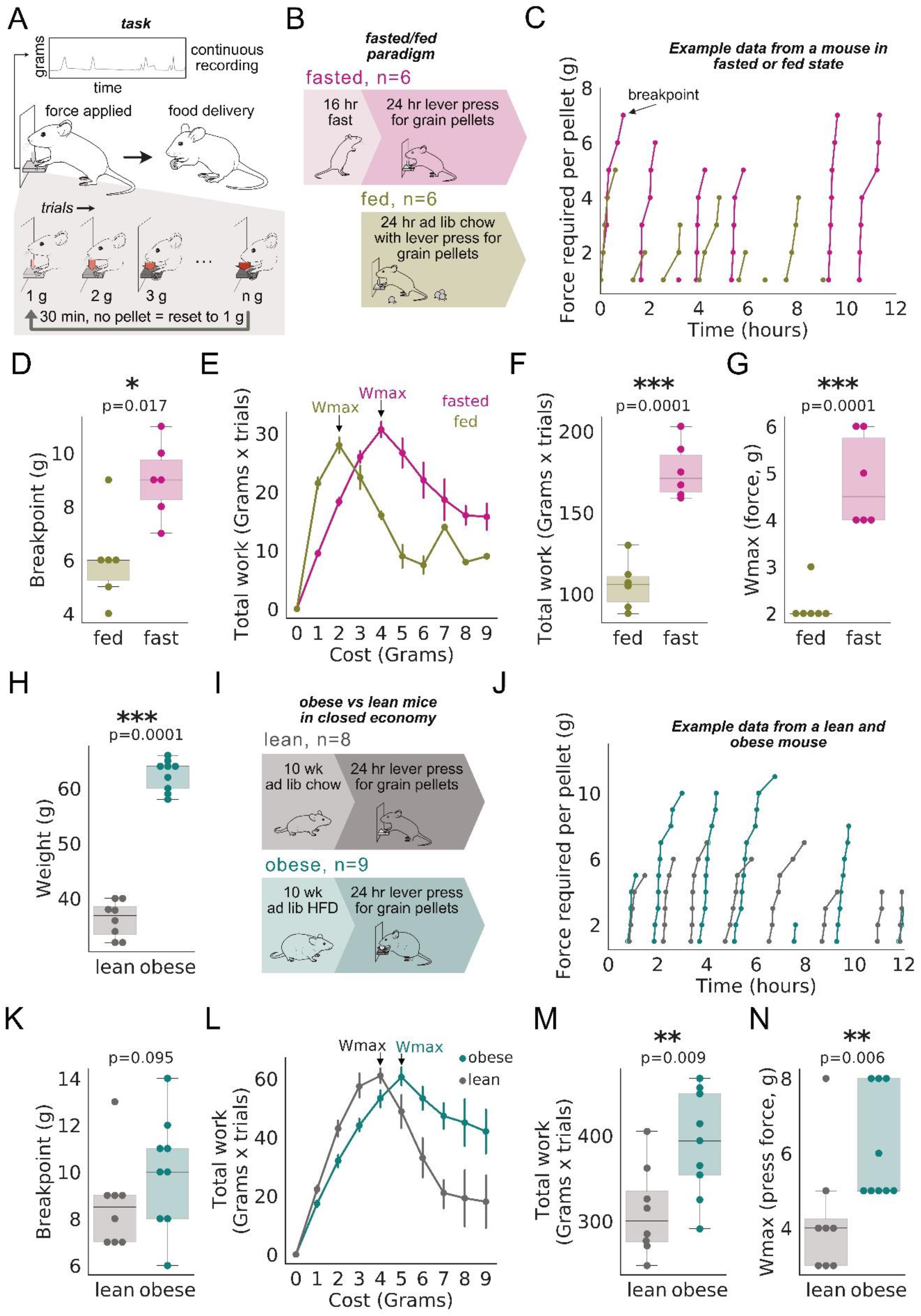
Obese mice work harder for food than lean mice. (a) Schematic illustrating closed economy food seeking task. Mice must press a lever to acquire 20 mg grain pellet. Subsequent trials require increased press force. Press force resets after 30 min without a successful trial. (b) Experimental setup: mice were fed ad lib chow or **HFD** for 10 weeks, trained on lever pressing task, and then exposed to closed economy task. (c) Example of force required per pellet in a resetting progressive ratio task from an example mouse in fasted (pink) and fed (olive) states. Each line represents a block of trials prior to reset. Breakpoint is the highest force achieved before reset.(d) Fasted mice exhibit higher max breakpoints than fed mice (p=0.017). (e) Work (required press force × number of trials completed) vs cost (required grams). (f) Total work (sum of work across all costs) increases in fasted mice relative to fed (p=0.0001). (g) Wmax (maximum work performed in a session) increases in fasted mice relative to fed, p =0.0001. (h) Mice exposed to ad lib **HFD** weigh more than lean mice (p=0.0001). (i)Experimental setup: mice were fed ad lib chow or HFD for 10 weeks, trained on lever pressing task, and then exposed to closed economy task. U) Force required per pellet in a resetting progressive ratio task from an example obese (teal) and lean (grey) mouse. Each line represents a block of trials prior to reset. (k) Max breakpoints between lean and obese, trend towards increase in obese, p=0.095. (I) Work (required press force × number of trials completed) vs cost (requ ired grams). (m) Total work (sum of work across all costs) increases in obese mice relative to lean, p=0.009. (n) Wmax (maximum work performed in a session) increases in obese mice relative to lean, p =0.006. Dependent ttest for within-subject comparisons in (d, f, g). Mann Whitney U test for comparisons in (h, k, m, n).

We first examined how differences in food motivation alter performance on this task by testing mice in either a fasted or fed state (Fig. 1B). Relative to the fed state, fasted mice reached significantly higher force breakpoints (Fig. 1D), performed significantly more total work (Fig. 1E,F), and had higher W_max_ values (the force requirement at which maximal total work was performed, Fig. 1G). Importantly, mice were slightly lighter in the fasted vs. fed state, yet exerted higher levels of force on the lever, demonstrating that lever-pressing force reflects changes in food motivation, and not changes in body weight. We next asked how performance of obese mice compared to lean mice on the same task (Fig 1H-N). Similar to fasted mice, obese mice trended towards higher force breakpoints (Fig 1J,K), performed more total work (Fig 1L,M), and had higher W_max_ values (Fig 1N) than lean mice. There were no significant correlations between body weight and work output in either lean or obese groups (Supp Fig 2A,B), demonstrating that, as in the fasted/fed experiment above, differences in work during food seeking were not related to the elevated body weights in obese mice. In summary, obese mice performed *more* physical work than lean mice to obtain food, clarifying that the relative reinforcing value of food in enhanced in obese mice, as it is in people with obesity^1,3,30^.

**Supplemental Figure 2.**
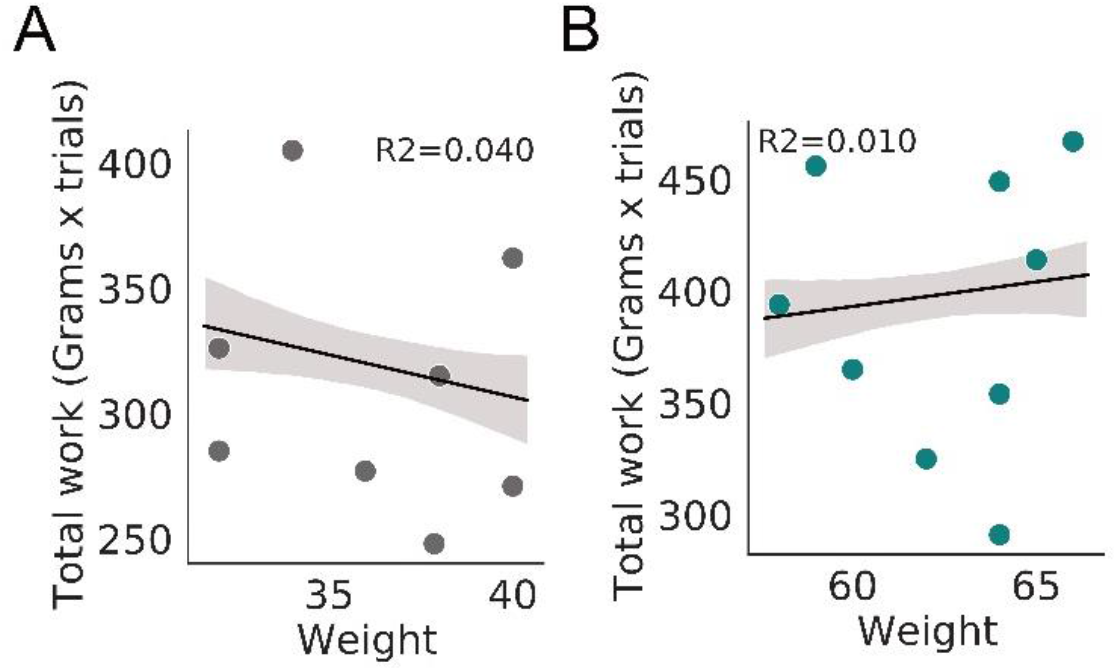
Body weight does not correlate with total work output or breakpoint in a closed economy food seeking task. Correlations between body weight and total work performed for lean (a) and obese (b) mice. Pearson correlations reveal no significant relationship between body weight and total work done or breakpoint in either group (R^2^=0.04 and 0.010 for lean and obese total work, respectively).

**Figure 2.**
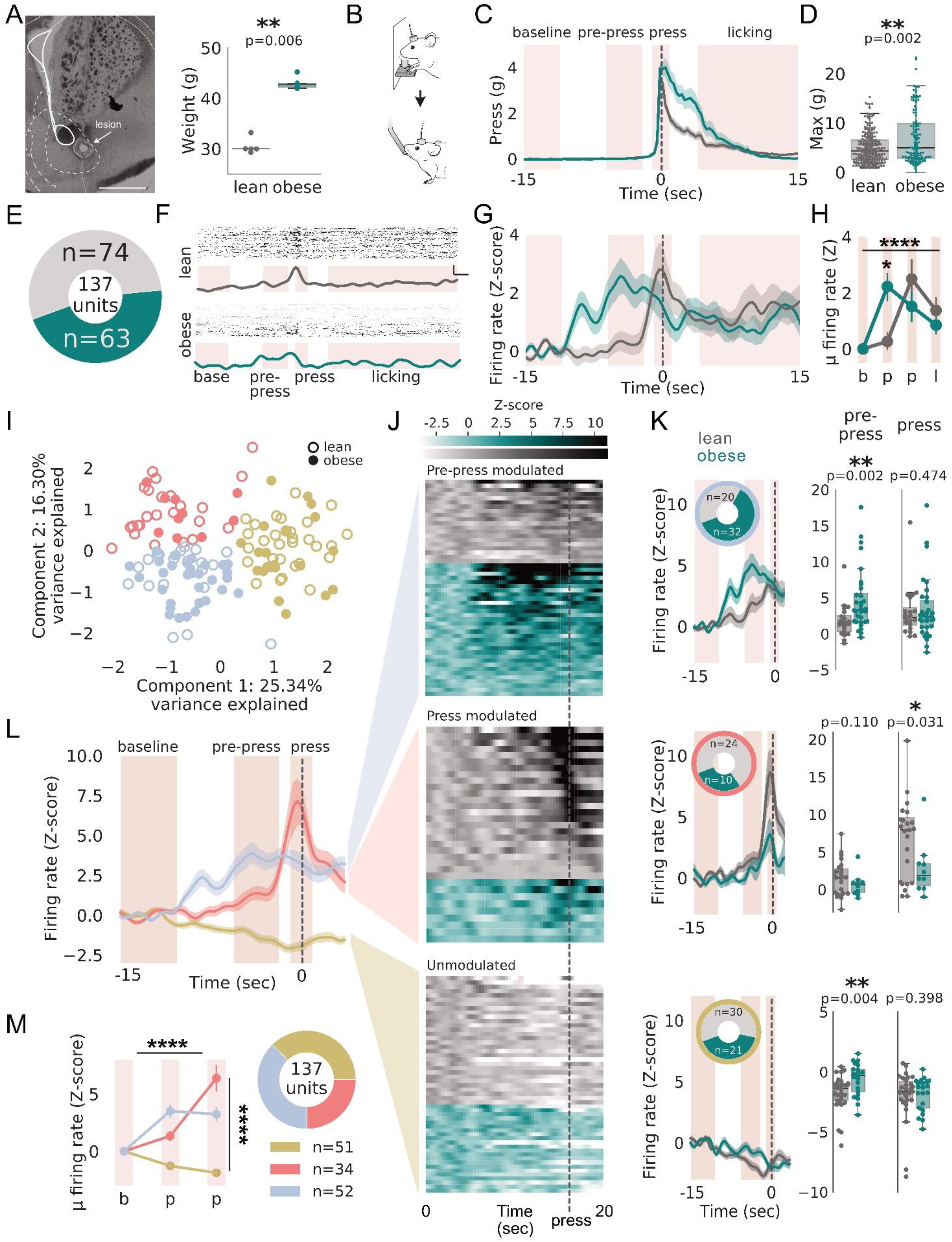
Obesity increases NAc firing during food seeking. (a) Left: Representative histological image showing Nae array implant location; scale bar: 1 mm. Right: Obese mice weigh significantly more than lean mice (p=0.006). (b) Mice with cranial implants press a lever with at least 2 g of force to recieve 20 uL of 5% sucrose. (c) Histogram of average lever press from lean and obese mice. Time O is reward delivery. (d) Max force per trial is significantly increased in obese mice relative to lean, p=0.002. (e) ring plot showing unit count for obese (n=63 units) and lean (n=74 units). (f) Example raster plots from each trial in a session and summary traces of a unit from lean (top, grey) and obese (bottom, teal) mice. Scale bar vertical=5 Z-scores, horizontal=2 sec. (g) Histogram of normalized firing frequency from lean and obese mice during food seeking. Time 0 is reward delivery. (h) Pointplot showing average unit firing rate normalized to baseline across pre-press, press, and lick periods. Significantinteraction between group and time (p=0.004, F(3,540)=4.508). Significant effect of time (p=0.0001, F(3,540)=7.607. No significant effect of group (p=0.706, F(1,540)=0.142). Tukey post-tests reveal sigficiant difference between groups during pre-press (p<0.05). (i) K-Means cluster analysis on principal components (PCs) 1, 2, and 3 identified three patterns of firing during food seeking. Scatter plot of PC1 vs PC2 for all units. U) Mean firing rate (z-scored relative to baseline) of all NAc units. Each row shows data for a single unit, sorted by group (lean, grey; obese, teal) and maximum value (descending) within each cluster. (k) Left: Histograms of clustered units separated by group (obese: teal, lean: grey). Inset: Ring plot showing number of units per group in each cluster. Right: Dotted bar graphs of average firing rates during food seeking phases, compared between lean and obese. Significant increase in firing frequency in obese during pre-press in blue and gold clusters. p=0.002, p=0.004, respectively. Significant decrease in firing frequency in obese during press in salmon cluster, p=0.031. No difference between groups for blue cluster press, salmon cluster pre-press, or gold cluster press (p>0.05 for all). (I) Histogram of clustered units during food seeking. (m) Average of clustered neural firing in baseline, b, pre-press, pO, and press, p1, for each cluster. Clustered firing patterns differ significantly: comparison of average firing in each period shows significant effect of cluster (F(2,402)=65.916, p=0.0001), food seeking phase (F(2,402)=17.069, p=0.0001), and interaction (F(4,402)=27.246, p=0.0001). Linear mixed model with multiple comparisons. Ring plot showing cluster unit counts. Mann whitney U test in (b,k). Dotted vertical line denotes press.

### Diet-induced obesity increases accumbal firing during food seeking

The NAc mediates behavioral reinforcement and controls vigor during food-reinforced tasks^32–36^. *Ex vivo* studies also suggest that NAc function is altered by exposure to obesogenic diets^14,16,37^. To test whether *in vivo* neuronal activity in the NAc was altered in obese mice during food seeking, we chronically implanted microwire arrays into the NAc of 5 lean and 4 obese mice (Fig. 2A) and recorded single- and multi-unit activity from these mice (74 units in lean mice, 63 units in obese mice). We first recorded each mouse for 60 minutes in an open field chamber, to determine if there were changes in the spontaneous firing properties of NAc neurons in obese mice. Firing rates in obese mice were lower relative to lean mice (Supp. Fig. 3A), and bursting activity in obese mice was characterized by longer bursts with fewer spikes per burst than lean mice (Supp. Fig. 3B-F). We next recorded each mouse during a task in which they had to perform a fixed cost 2-gram press to obtain a 20 μL drop of 5% sucrose solution (Fig. 2B-D). Similar to the increases in physical work we observed in the progressive force task (Fig. 1), obese mice exerted higher peak forces and held these forces longer than lean mice did, despite there being no incentive to do so (Fig. 2B-D), which we observed in a separate cohort of lean and obese mice (Supp. Fig. 4). In lean mice, average NAc unit activity increased sharply at ∼500ms before the lever press (Fig. 2E-H), consistent with prior reports of NAc firing patterns during reward seeking^38,39^. In obese mice, this response was shifted earlier in time, with firing increasing for several seconds prior to the lever press (Fig. 2E-H). This is consistent with the change in bursting activity we observed (Supp Fig 3), in which bursts were longer but of lower frequency in lean mice.

**Figure 3.**
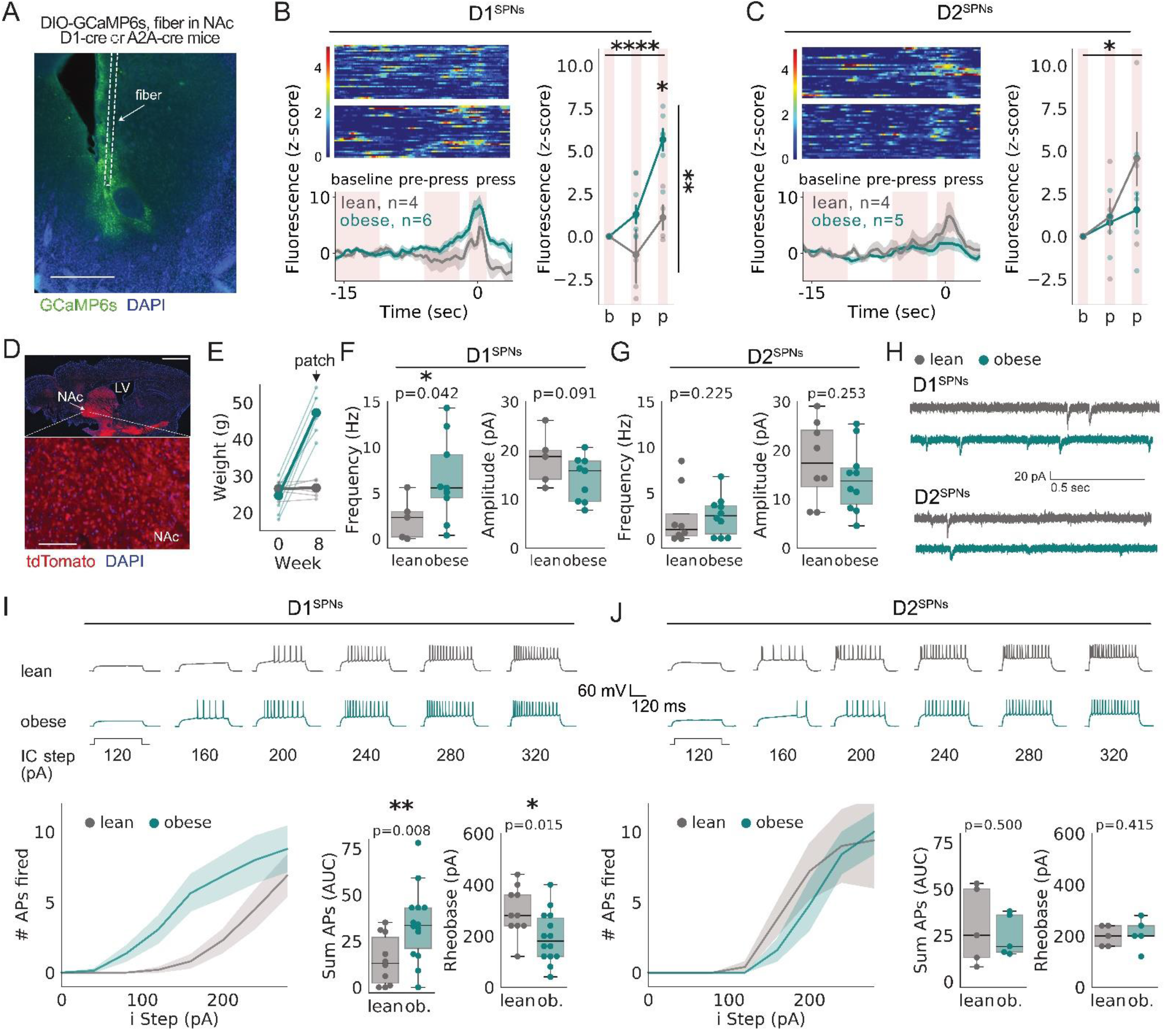
Obesity increases excitation of NAc D1^sPNs^. (a) Schematic showing DIO-GCaMP6s injection and fiber implant in NAc core of 01-Cre or A2A-Cre mice. Representative histological image. Scale bar: 1 mm. (b) Heat maps showing every trial from an example food seeking session. Bottom: Perievent histogram plotting z-scored GCaMP6s fluorescence from lean and obese 01- Cre mice. Right: Pointplot with averaged fluorescence showing significant increase in fluorescence in obese D1sPNs relative to lean (Significant effect of time (p=0.0001, F(2,24)=16.184), group (p=0.001, F(1,24)=4.493), and interaction between group and time (p=0.0004, F(9,111)=2.907). Tukey post hoc tests reveal significant difference between lean and obese during press (p<0.05)). (c) Heat maps showing every trial from an example food seeking session. Bottom: Perievent histogram plotting z-scored GCaMP6s fluorescence from lean and obese A2A-Cre mice. Right: Pointplot with averaged fluorescence showing no significant change in fluorescence in obese D2^sPNs^ relative to lean (Significant effect of time (p=0.0467, F(2,21)=3.559. No significant effect of group (p=0.231, F(1,21)=1.523), nor interaction between group and time (p=0.348, F(2,21)=1.110). (d) Acute striatal sections were prepared from 01-tomato mice; labelled (putative 01) and unlabelled (putative 02) SPNs were patched in NAc core in lean and obese mice. Histological image of parasaggital section of 01-tdTomato mouse stained with DAPI (blue). Tomato is red. LV: lateral ventricle. Top, scale bar is 2 mm, bottom, scale bar is 250 um. (e) Weights over time as mice gained weight (obese group, teal) or were maintained on chow (lean, grey). (f) Frequency of mEPSCs increases in NAc D1-SPNs in obese mice vs lean (p=0.042); no significant difference in amplitude of mEPSCs in D1^sPNs^ between lean and obese (p=0.091). n = 5(3) lean, 9(4) obese. (g) No significant difference in D2-SPN mEPSC frequencey (p=0.225), or amplitude (p=0.253) between groups. n = 9(4) lean, 11(4) obese. (h) Example traces from lean (grey) and obese (teal) D1- (top) and D2- (bottom) SPNs. (i) Example traces and excitability curves of evoked firing in D1^sPNs^ from lean and obese mice. Significant increase in area under the curve (AUG, summation of APs) and significant decrease in rheobase (pA) in obese 01-SPNs relative to lean (p=0.008, p=0.005, respectively). n= 10(6) lean,14(4) obese. 0) Example traces and excitability curves of evoked firing in D1^sPNs^ PNsfrom lean and obese mice. No signficiant difference in AUC or rheobase in obese D2^sPNs^ relative to lean (p=0.464, p=0.463, respectively). n = 5(5) lean, 5(4) obese. Mann Whitney U test for comparisons in (b, g, h, j, k).

**Figure 4.**
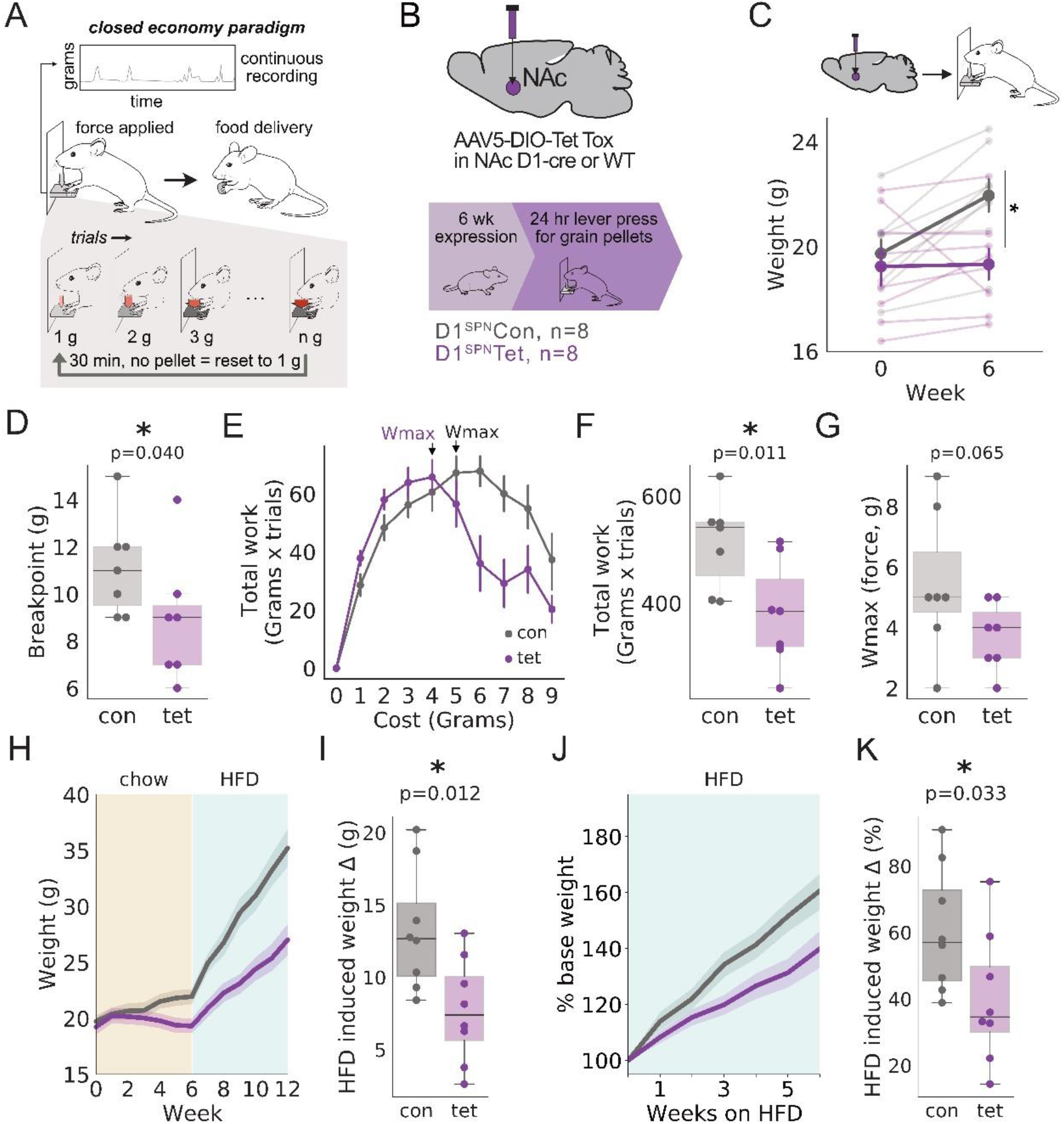
Blocking D1^sPNs^ decreases food motivation and prevents obesity. (a) Schematic illustrating closed economy food seeking task. Mice must press a lever to acquire 20 mg grain pellet. Subsequent trials require increased press force. Press force resets afte 30 min without a successful trial.) (b) D1-cre (tet) or WT (con) mice were injected with DI0-tet in NAc core and virus was allowed to express for 6 weeks before closed economy task. (c) Pointplots showing weight change following cranial injections (Week 0). Food seeking assay was conducted at Week 7 post-surgery. Significant effect of group (p=0.0268, F(1,28)=5.464). No significant effect of time (p=0.097, F(1,28)=2.941) or interaction between group and time (p=0.122, F(1,28)=2.539). (d) Breakpoint decreased in tet-tox mice relative to control (p=0.040). (e) Work (required press force × number of trials completed) vs cost (required grams). (f) Total work (sum of work across all costs) decreases in tet-tox mice relative to control, p=0.011. (g) Wmax (maximum work performed in a session) between control and tet-tox, p =0.065. (h) Con and tet mice were exposed to ad lib chow in the homecage for seven weeks. At week six mice were given ad lib HFD for six weeks. Mice were weighed weekly. Weights over time for con (grey) and tet (purple) groups. (i) Tet mice gained significantly less weight than con, p=0.012. (j) Percent change in weight over time after ad lib HFD. (k) Tet mice gained signifcantly smaller percentage of body weight from start of HFD than con, p=0.033. Mann Whitney U test for comparisons in (d, f, g, i, k). Mixed linear model with repeated measures in (c).

NAc SPNs can encode distinct elements of a behavior such that they fire selectively at different phases of a behavioral sequence^40–42^. To determine whether changes in NAc firing during food seeking in obese mice reflected a specific response type, we classified neural firing patterns during the lever press. A K-means clustering algorithm revealed three distinct motifs (Fig. 2I, L, M), including units that: 1) increased firing during the pre-press period (blue cluster), 2) increased firing during the lever press itself (salmon cluster), or 3) were not activated during either of these periods (gold cluster). Consistent with what we observed at the population level, units in the pre-press cluster were significantly more likely to come from obese than lean mice (Supp Fig. 5A). Average firing rates of units in the pre-press cluster from obese mice fired at significantly higher rates than those from lean mice (Fig. 2J,K top). Units modulated during the lever press (salmon cluster) were predominantly found in lean mice (Supp Fig. 5B), and reached higher firing rates during the lever-press than those from obese mice (Fig. 2J,K middle). Finally, lean mice contributed more units than obese mice to the “non-activated” cluster though this was not significantly different from chance (Supp Fig. 5C) and these units displayed lower firing frequencies during the pre-press period than those contributed from obese mice (Fig 2J,K bottom). We conclude that NAc activity during food seeking is disordered in obese mice, with the majority of neural firing occurring earlier – and firing significantly faster – in the press sequence relative to lean.

**Supplemental Figure 3.**
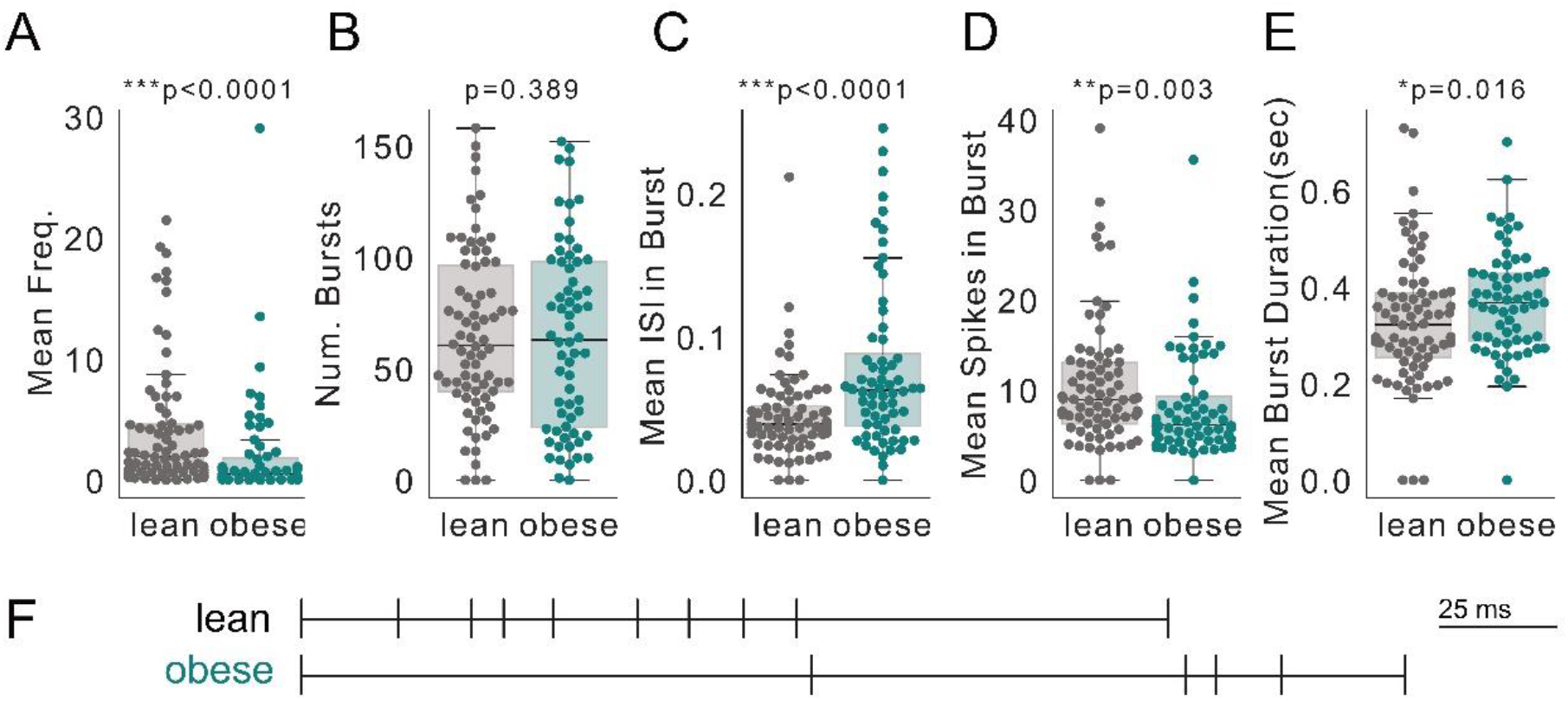
Obesity decreases basal NAc firing frequency. (a) Average firing frequency of units recorded in obese mice was significantly lower than those recored in lean, p=0.0001. (b) No significant difference in number of bursts identified in lean vs obese mice, p=0.389. Burst analysis revealed significant increase in inter-spike interval (ISi) (c), significant decrease in average number of spikes per burst (d) and significant increase in average burst length (e) in obese vs lean. p=0.0001, 0.003, and 0.016, respectively. (f) Schemative illustrating longer-duration burst firing with fewer spikes per burst in obese. Mann Whitney U test in a-e.

**Supplemental Figure 4.**
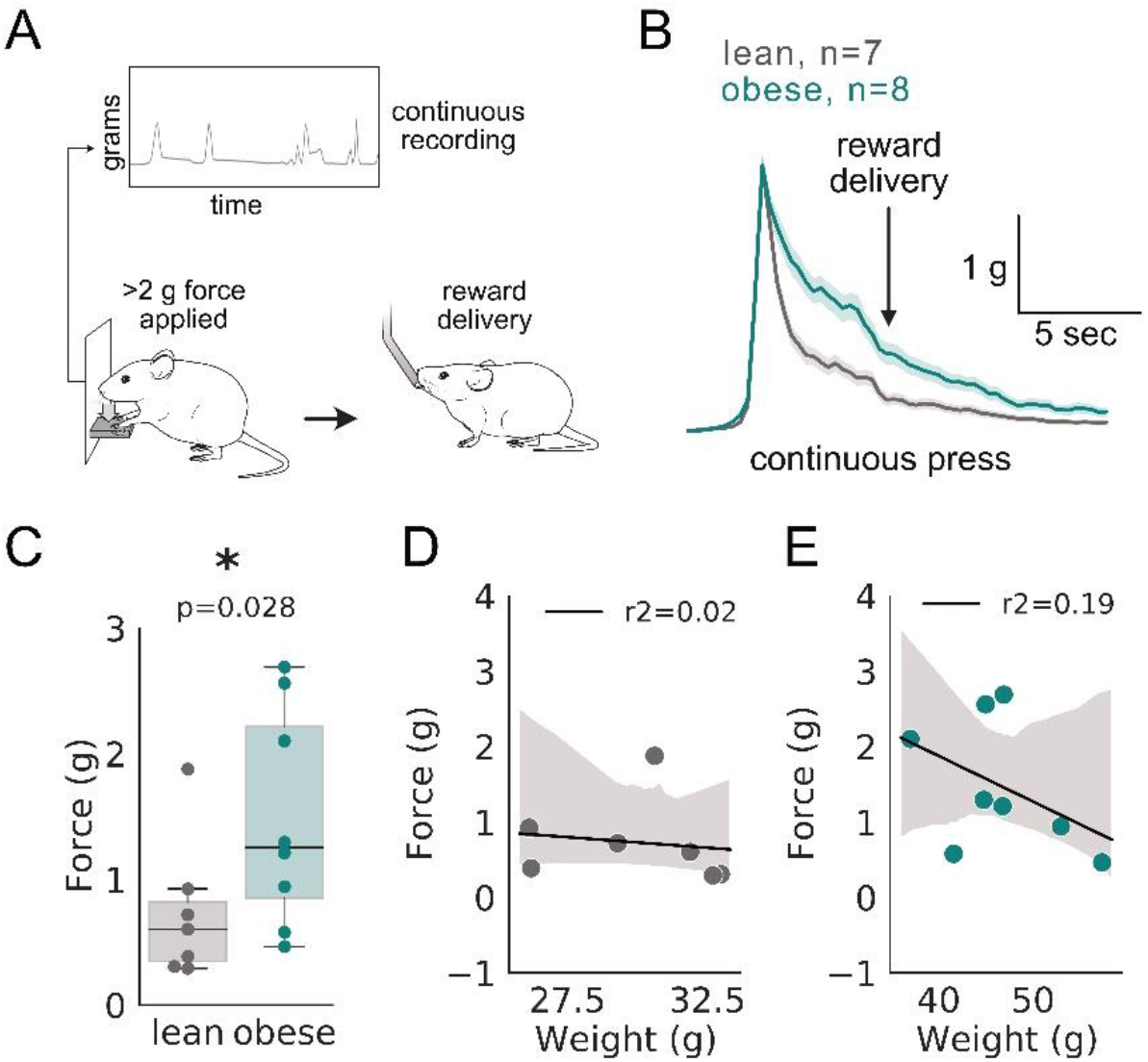
Obesity increases effort during food seeking. (a) Schematic showing fixed ratio lever pressing tas. Mice learn to press a lever with at least 2 g of force to recI ve 20 ul of 5% sucrose. (b) Continuous lever press profile from lean and obese mice. (c) Average force output per mouse was signficiantly higher in obese mice relative to elan, p=0.028. (d,e) Correlations between body weight and force output performed for lean (d) and obese (e) mice. Pearson correlat1<:;>ns reveal no significant relationship between body weight and force output in either group (R2=0.00 and 0.15 for lean and obese respectively.) Mann Whitney U test in c.

**Supplemental Figure 5.**
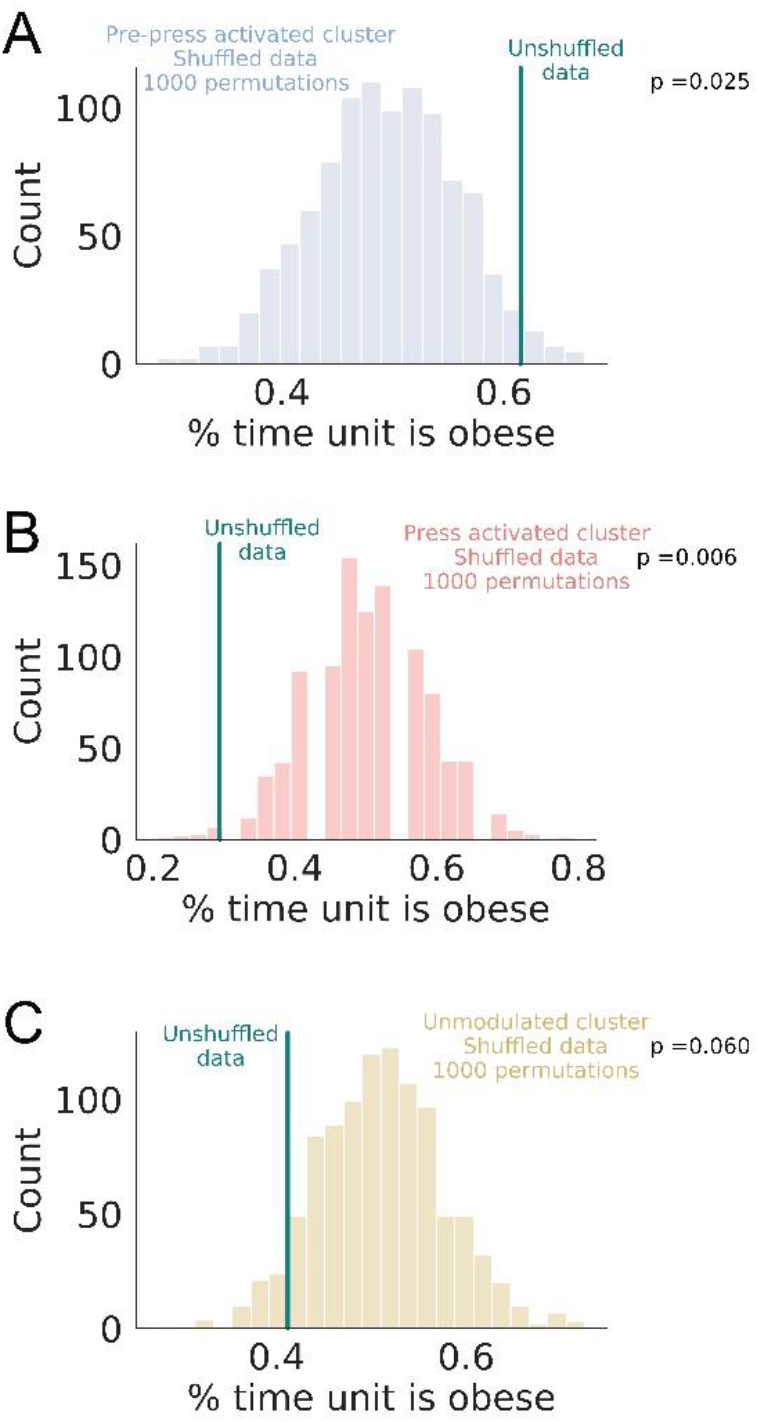
Permutation analyses for clustered spike patterns. Distributions of average percent obese units between obese and lean categorizations with 1,000 permutations of shuffled categorizations for (a) pre-press modulated, (b) press-modulated, and (c) unmodullated clusters. Teal vertical lines are ground truth percent obese per cluster. Z-tests reveal obese unit percentage is significantly different from a random distribution at the p= 0.025 (a), p= 0.006 (b), and p= 0.060(c) level.

### Obesity preferentially enhances excitatory input to NAc D1-SPNs

We next asked whether NAc D1^SPNs^ or D2^SPNs^ contributed to the increased activity observed in our electrophysiological recordings. We expressed a Cre-dependent genetically encoded calcium indicator (GCaMP6s) and implanted fiber optics in the NAc core of lean and obese D1-Cre or A2A-Cre transgenic mice, to target expression to D1^SPNs^ and D2^SPNs^ respectively (Fig. 3A). Mice performed the same task described in Figure 2 while we recorded bulk calcium activity from NAc D1^SPNs^ and D2^SPNs^ (Fig. 3B-C). Population calcium activity of D1^SPNs^ was enhanced during the pre-press and press period in obese mice relative to lean mice (Fig. 3B), consistent with our previous electrophysiological recordings. In contrast, we observed no differences in D2^SPN^ calcium activity in these same periods (Fig. 3C). This suggests that increased NAc spiking activity during food seeking in obese mice was specific to D1^SPNs^.

Recordings of bulk calcium activity can include subthreshold fluctuations in both dendritic and somatic calcium^43^. To test whether increases in D1^SPN^ calcium activity reflected increased excitatory drive onto D1^SPNs^, we performed whole cell recordings of spontaneous excitatory events in the presence of tetrodotoxin (TTX), which allowed us to record quantal release in the absence of action potential-mediated vesicular release (mini excitatory post-synaptic currents, or mEPSCs). In D1-tomato transgenic mice, we patched visually identified (D1^SPN^) and unlabeled (putative D2^SPNs^) in the NAc core (Fig. 3D) of lean or obese mice (Fig. 3E). Consistent with our fiber photometry results, we observed a higher frequency of mEPSCs onto D1^SPNs^ but not D2^SPNs^ of obese mice relative to lean mice (Fig. 3F-H)). No significant changes were observed in mEPSC amplitude in either cell type between groups (Fig. 3F-H). This supports the conclusion that obesity selectively facilitates excitatory drive onto D1^SPNs,^ via increasing the presynaptic probability of release.

Prior work has also implicated alterations in cell-autonomous NAc function in obese animals. Specifically, exposure to obesogenic diets has been shown to increases in intrinsic excitability of NAc neurons^14,15^, although it has not been determined whether these changes occur specifically in D1^SPNs^ or D2^SPNs^. To test this, we prepared acute striatal sections from lean and obese mice and evoked action potential firing in identified D1^SPNs^ and D2^SPNs^ with successively increasing current steps. We observed an increase in the current-evoked spiking of D1^SPNs^ but not D2^SPNs^ in obese mice relative to lean mice (Fig 3I-J). Further supporting this dissociation, injections of ramp currents revealed a decrease in rheobase in D1^SPNs^ but not D2^SPNs^ from obese mice relative to lean mice (Fig 3I-J). Additionally, there were no differences between lean and obese D1^SPN^ or D2^SPN^ resting membrane potential (lean D1, μ = −71.4 +−6.418 mV, obese D1, μ = −68.62 +− 6.784 mV, Student’s t-test, p=0.163; lean D2, μ = −69.9 +− 7.651 mV; obese D2, μ = −73.6 +− 2.966 mV; Student’s t-test, p=0.071.). Together our *ex vivo* slice studies demonstrate both pre- and post-synaptic effects that increase D1^SPN^ signaling in obese mice, consistent with the enhanced *in vivo* D1^SPN^ activity observed during food seeking.

To test our hypothesis that the balance of D1^SPN^:D2^SPN^ activity was shifted towards D1^SPNs^ in obese mice, we re-analyzed population-specific photometry and cell-type-specific patch clamp data within each group of lean or obese mice. While there were no significant differences in D1^SPN^ vs. D2^SPN^ calcium activity around the lever-press of lean mice, D1^SPNs^ were significantly more active than D2^SPNs^ in obese mice, with significantly higher fluorescence during press (Supp Fig. 6A,B). Similarly, there was no significant difference in the frequency or amplitude of mEPSCs between D1^SPN^ and D2^SPN^ in lean mice, but a significantly higher frequency (but not amplitude) of mEPSCs in D1^SPNs^ relative to D2^SPNs^ in obese mice (Supp Fig 6C,D). Finally, in lean mice, we observed a trend towards increased intrinsic excitability and a significant decrease in rheobase in D2^SPNs^ relative to D1^SPNs^, as previously reported^44^(Supp Fig 6E). However, this difference in excitability was no longer apparent in obese mice. Thus, we observed a shift in the balance of activity towards D1^SPN^ over D2^SPN^ signaling in obese mice.

**Supplemental Figure 6.**
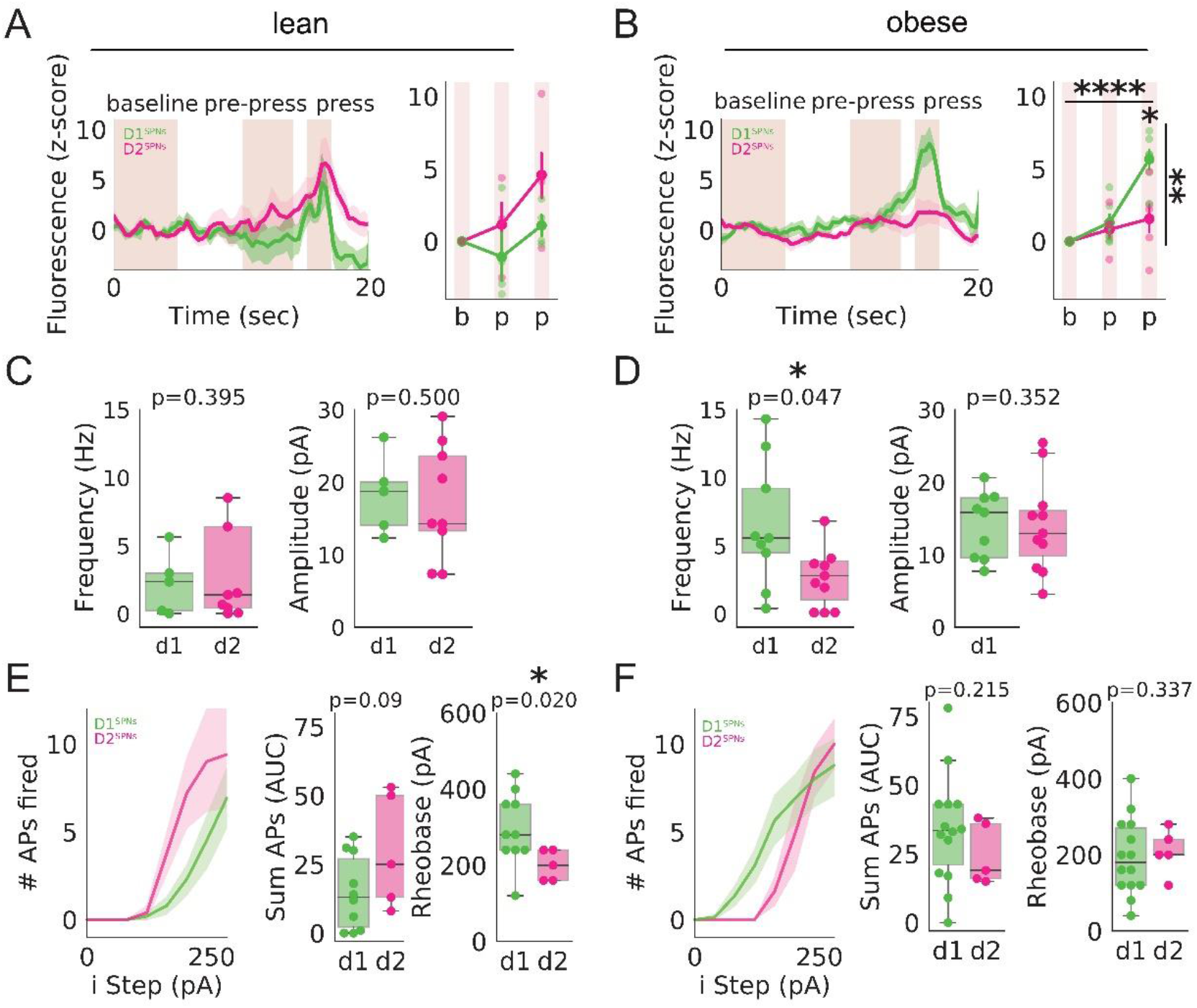
Obesity skews NAc D1/D2 balance, (a) Left: Perievent histogram plotting z-scored GCaMP6s fluorescence from lean D1-Cre and A2A-Cre mice. Right: Pointplot with averaged fluorescence showing significant increase in fluorescence in obese D1-SPNs relative to D2 (No significant effect of time (p=0.066, F(2,18)=3.164), genotype (p=0.089, F(1,18)=3239), nor interaction between genotype and time (p=0.413, F(2,18)=0.928). (b) Left: Penevent histogram plotting z-scored GCaMP6s fluorescence from obese D1-Cre and A2A-Cre mice. Right: Pointplot with averaged fluorescence showing significant increase in fluorescence in obese D1-SPNs relative to D2. (Significant effect of time (p=0.0001, F(2,27)=18.314, and genotype (p=0.008, F(1,27=8.317), and interaction between genotype and time (p=0.007, F(2,27)=6.012). Tukey post hoc tests reveal significant difference between D1 and D2 during press (p<0.05). (c) Frequency of mEPSCs in NAc D1-SPNs and D2-SPNs in lean mice (p=0.395); amplitude of mEPSCs in D1-SPNs and D2-SPNs in lean mice (p=0.500). (d) Frequency of mEPSCs in NAc D1-SPNs is significantly higher than D2-SPNs in obese mice (p=0.047); amplitude of mEPSCs in D1-SPNs and D2-SPNs in obese mice (p=0.352). e) No significant difference in area under the curve (AUC, summation of APs) between cell types in lean mice; significant decrease in rheobase (pA) of D2-SPNs in lean (p=0.058, p=0.014, respectively), (f) No significant difference in AUC or rheobase between cell types in obese mice (p=0.215, p=0.406, respectively). Linear mixed model in (a, b). Mann Whitney U test for comparisons in (c-f).

### Silencing NAc D1^SPNs^ decreases the vigor of food seeking and attenuates diet-induced obesity

Our results thus far indicate that food seeking vigor is enhanced in obese mice, who also exhibit a selective increase in D1^SPN^ activity during food seeking. Therefore, we explicitly tested whether D1^SPNs^ were necessary for invigorating food seeking. To accomplish this, we silenced the output of D1^SPNs^ by virally expressing a Cre-dependent tetanus toxin subunit (CBA-DIO-GFP-Tet Tox-WPRE-bHGpA) in the NAc of D1-Cre-positive (D1^SPN^Tet) or -negative littermate (D1^SPN^Con) mice (Fig 4B). Gross locomotor measures in the open field were not affected by the viral manipulation (Supp Fig 7). However, D1^SPN^Tet mice exerted less effort than D1^SPN^Con mice on the progressive force task (Fig 4A,C), as evidenced by significantly lower breakpoints, total work, and Wmax values (Fig 4. D-G). Consistent with prior reports^45^, we conclude that inhibiting the output of D1^SPNs^ does not produce a generalized motor deficit, but specifically reduces behavioral vigor during food seeking.

No weight differences were present between D1^SPN^Tet and D1^SPN^Con prior to surgery. Following surgery, all mice were maintained on lab chow for 6 weeks. At 6 weeks, D1^SPN^Tet mice were slightly but significantly lighter than D1^SPN^Con mice (Fig 4C). Based on their slightly lean phenotype, we tested whether D1^SPN^Tet mice would also be resistant to high-fat diet induced weight gain. We maintained both groups of mice on *ad libitum* high fat diet for six weeks (Fig 4H,J), and found that D1^SPN^Tet mice were resistant to gaining weight, relative to D1^SPN^Con mice (Fig 4H-K). Even when normalized to their body weights when high-fat diet was first introduced, D1^SPN^Tet mice had a significantly slower rate of weight gain than D1^SPN^Con mice (Fig 4J,K). Neither initial body weight nor HFD-induced weight changes correlated with work output during food seeking (Supp Fig. 8), demonstrating that decreases in work during food seeking were not due to decreased body weight in D1^SPN^Tet mice. We conclude that blocking D1^SPN^ output reduces both food seeking vigor and high-fat diet induced weight gain.

**Supplemental Figure 7.**
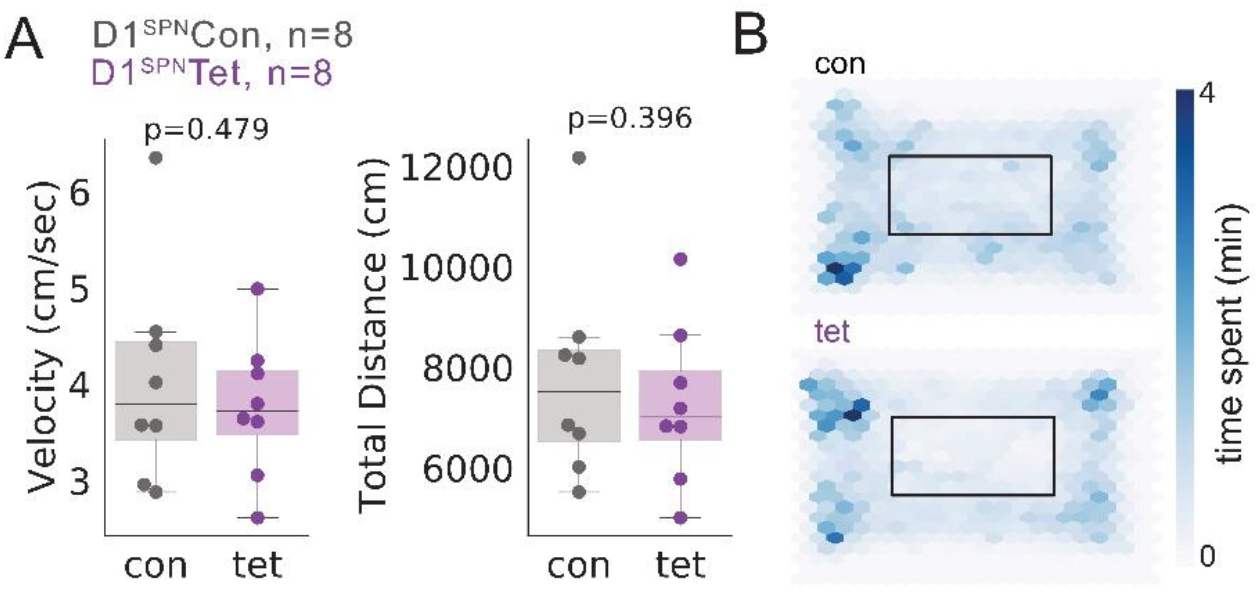
Blocking D1^sPNs^ does not change average velocity during an open field experiment (a) Average velocity and distance do not differ between control and tet-tox mice, p=0.479 and p=0.396, respectively. Mann Whitney U test. (b) Compiled locomotion heat maps showing control (top) and tet-tox (bottom) mouse location in open field arenas. Scale bar, right, showing relative time spent across the arena.

**Supplemental Figure 8.**
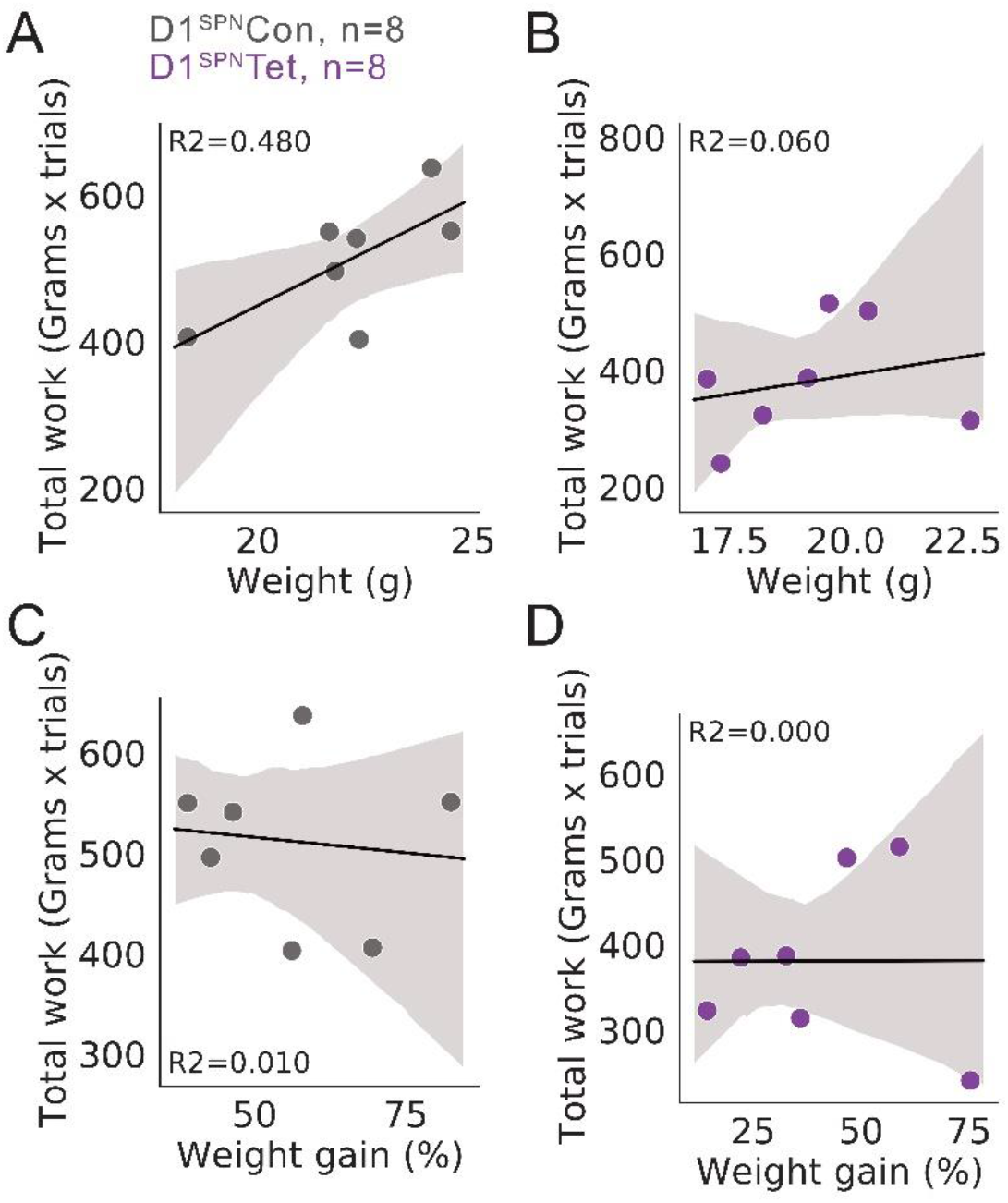
Work output for food does not correlate with weight or weight gain after **HFD**. Correlation between body weight (week 7) and total work performed for control (a) or tet tox (b)mice. Pearson correlation revealed no significant relationship between body weight and total work done (R^2^=0.07).

## Discussion

Obesity has been linked to increases in the relative reinforcing value of food^4,5^, which can lead to overeating and weight gain^1,3^. However, the neural circuits that underlie changes in food reinforcement in obese animals remain unknown. Here, we tested the hypothesis that obesity-linked increases in the relative reinforcing value of food are driven by changes in the NAc core^46–48^, shifting the balance of activity towards D1^SPNs^ over D2^SPNs^ during food seeking. Consistent with this hypothesis, NAc D1^SPN^, but not D2^SPN^, activity was enhanced in obese mice during food seeking, which was linked to both pre- and post-synaptic adaptations in D1^SPNs^ in an *ex vivo* preparation. Blocking synaptic output from D1^SPNs^ reduced food seeking vigor and attenuated high-fat diet induced weight gain. This provides a circuit-based explanation for changes in food reward and reinforcement in people with obesity, and highlights NAc D1^SPNs^ as a potential therapeutic target for treating or preventing obesity.

NAc D1^SPNs^ have been linked to both behavioral reinforcement and the invigoration of ongoing motor actions^19,20^. Therefore, we hypothesized that an increase in the relative reinforcing value of food may be driven by increased D1^SPN^ activity during food seeking in obese mice. Excitatory adaptations have been previously reported in NAc core SPNs in obesity prone rats^15,49^, but it was not clear from prior research whether such changes were selective for D1^SPNs^ or D2^SPNs^. Here, we found that excitatory adaptations in obese mice occurred selectively in D1^SPNs^, supporting work showing that NAc pharmacological inhibition or global knockout of D1 receptors attenuates food seeking^50,51^ and prevents obesity in rodents^51^.

Obesity has been linked to post-synaptic insertion of calcium-permeable alpha-amino-3-hydroxy-5-methylisoxazole-4-propionic acid (AMPA) receptors in NAc core^14^ – a process that facilitates early-phase long-term potentiation (LTP) induction^52^ and can underlie enhancements in food motivation^53^. Based on this, we expected to observe changes in post-synaptic mEPSC amplitude, which we did not. However, we observed enhancements in intrinsic excitability and *pre-synaptic* drive, both selectively in D1^SPNs^. These may reflect additional synaptic adaptations that are consistent with disruptions in long-term depression (LTD) induction, which has been observed *ex vivo* in NAc SPNs from obesity-prone rats^16^. Thus, while obesity likely drives maladaptive plasticity in the NAc, further work is needed to determine the locus of effect and exact mechanisms affected.

In contrast to our present findings, prior literature has also demonstrated a role for NAc D1^SPNs^ in inhibiting food consumption. Specifically, optogenetic activation of NAc medial shell D1^SPN^ projections to the lateral hypothalamus^54^ or ventral tegmental area^55^ halts ongoing consumption of food. Given that we observed enhanced activity and excitability of NAc D1^SPNs^ in obese mice, shouldn’t these adaptations lead to an inhibition of food consumption? The answer to this question arises from the specific phases of feeding being studied and from the anatomic specificity of NAc subregions that mediate each phase. Feeding includes temporally distinct phases of food seeking, food consumption, and food evaluation^56^. Whereas D1^SPN^ activity increases during food seeking, most D1^SPNs^ (and most D2^SPNs^) are inhibited during consumption itself^57^. Prior work suggesting that D1^SPNs^ inhibit feeding is consistent with D1^SPN^ projections acting as a “sensory sentinel” to rapidly stop feeding after it starts, if spoiled food or other danger is detected^58^. This sensory sentinel function is not likely to be engaged during consumption of palatable foods in our assay.

In addition, the specific NAc subregion is critical for interpreting the effect of D1^SPN^ manipulations on food seeking and consumption. Manipulations in the NAc medial shell (the region targeted in the optogenetic experiments above^54^) have been linked most strongly to changes in food consumption, while manipulations in the NAc core (the region targeted in our present experiments) have been linked most strongly to changes in food seeking and behavioral vigor. For example, infusions of the AMPA receptor antagonist 6,7-dinitroquinoxaline-2,3-dione (DNQX) into the NAc medial shell, (but not the NAc core) increased food consumption^59,60^. Inhibiting the NAc shell by infusing the gamma-aminobutyric acid (GABA)-A receptor agonist muscimol produced similar effects, suggesting that overall activity levels within the NAc shell are sufficient to control consumption^61,62^. In contrast, manipulations of activity in the NAc core tends to not alter food consumption, but rather alter the expression and vigor of food seeking. For example, infusion of GABA agonists in core or shell diminished or enhanced responding to a learned food cue, respectively, in rats^63^, and infusion of the NMDA antagonist AP-5 into the core (but not shell) blocked acquisition of a bar press to receive food in rats^64^. Our experiments focused on the NAc core and are consistent with the appetitive role of D1^SPNs^ in behavioral vigor and food seeking.

One important prediction arising from this work is that inhibition of NAc D1^SPNs^ may be a therapeutic target for promoting weight loss or preventing weight gain. In rodents, global removal of D1Rs, or a selective removal of D1Rs in GABAergic neurons throughout the brain blocked high-fat diet induced weight gain^51^. The efficacy of a systemic D1/D5 antagonist ecopipam has also been tested in a clinical trial for weight loss in people with obesity, resulting in promising levels of weight loss^65^. However, this trial was terminated in Phase 3 due to the emergence of psychiatric side effects. This is not surprising, as ecopipam is not selective for SPNs, and targets dopamine receptors throughout the brain. It therefore remains possible that more targeted approaches acting on NAc core D1^SPNs^, or other selective subsets of D1^SPNs66^ may be able to modify body weight without adverse psychiatric effects. Our findings would predict selective inhibition of NAc D1^SPNs^ would reduce the vigor of food seeking and food reinforcement, and thereby cause weight loss. Together, our findings establish a model integrating adaptations in D1^SPNs^ with enhanced food seeking and weight gain. These findings lay the foundation for further investigations aimed at circuit-specific manipulation of NAc D1^SPNs^ to promote weight loss.

## Methods

### Mice

Adult male and female mice (12-20 weeks) were bred in-house on a C57Bl6 background. Transgenic lines were obtained from Jackson Laboratories (JAX) (D1-To, B6.Cg-Tg(Drd1a-tdTomato)6Calak/J; wild type C57Bl6/J) or the GENSAT project (A2a-Cre, Tg(Adora2a-cre)KG139Gsat; D1-Cre, Tg(Drd1-cre)EY217Gsat)^67^ and bred with wild type mice. Littermate controls were Cre negative littermates from these lines. Genotypes were confirmed by PCR and sequencing. All animal procedures were approved by the NIDDK or Washington University School of Medicine’s Institutional Animal Care and Use Committee and conformed to the National Institutes of Health guidelines. Mice were group housed (2-5 mice per cage) unless otherwise noted. Standard chow diet and water were given ad libitum. For experiments where obesity was induced, 60% high fat diet (Research Diets, D12492) was given ad libitum. No mice had drug exposures before or after stereotaxic surgery, and all mice were of normal immune status. Mice were housed in a 12 hour light/dark cycle in rooms that were controlled for temperature and humidity.

### Behavioral assays

#### Closed economy food seeking

Operant behavior was tested with the FORCE behavioral device, a custom operant device that senses weights applied to active and inactive isometric levers in a task dispensing food pellets (https://hackaday.io/project/163396-force-force-output-of-rodent-calibrated-effort). A lever press on the active lever yielded a combined tone and 20 mg grain pellet (TestDiet, 5TUM). Effort requirement started at 1 gram and increased by 1 g with subsequent trials. 30 minutes without a successful trial yielded a reset to a 1 gram requirement. All food available to mice was earned through lever pressing unless otherwise stated. Mice were run for 48 hour sessions, with ad libitum water available throughout.

#### Fixed ratio operant behavior

Operant responding was also tested with the FORCE device, although here it was connected to a solenoid to dispense liquid rewards. Tasks were conducted on a fixed ratio (FR) schedule, where sated mice learned to press an isometric lever with >2 grams. One successful press on the active lever yielded a tone and 20 uL of sucrose water (5%) and was immediately followed by a 30 second timeout. Mice were run for 16 hour sessions during the dark cycle and had ad libitum food available in the arena. Behavioral data was collected concurrently with electrophysiology.

#### Open field test

Mice were placed in the center of an arena (17 × 31 × 25 cm^3^) and locomotion was recorded with a usb-powered webcam for 30 minutes. The open-source Bonsai programming language^68^ was used to calculate x- and y-positions, and distance travelled was determined using Pythagorean Theorem. Velocity was calculated by dividing distance travelled by exact time in the arena.

### Stereotaxic surgery

Mice were deeply anesthetized with isoflurane (induction: 2-3%, maintenance: 0.5-1.0%) and secured in a stereotaxic frame (David Kopf Instruments), and craniotomies were performed. Purified AAVs were injected into the NAc core (relative to Bregma: anteroposterior (AP), +1.5 mm; mediolateral (ML), +/- 1.0 mm; dorsoventral (DV), − 3.85 mm). 300 nL of virus diluted to ∼10^12^ viral particles per mL was injected bilaterally into the NAc core. GCaMP6s virus (AAV-DJ-Ef1a-DIO-GCaMP6s) was obtained from Addgene, Tet-tox virus (AAV-CBA-DIO-GFP-TetTox-WPRE-bHGpA, plasmid was a gift from the Palmiter Lab^69^) was obtained from the Washington University Hope Center Viral Vectors Core. Injections used a pulled glass micropipette and delivered virus at a rate of 0.05 uL/min. After infusion pipettes were left *in situ* for an additional 10 minutes before being slowly withdrawn to allow for diffusion. For *in vivo* electrophysiology experiments, 16- or 32-channel tungsten microwire arrays (35 microns diameter, Innovative Neurophysiology) were lowered slowly (0.1 mm/min) and implanted into the NAc. For fiber photometry, optical fibers (fiber core: 200 micron core, 0.48 NA, ThorLabs) were implanted into the NAc. Arrays or fibers were unilaterally cemented onto the skull with a layer of dental cement (C&B Metabond; Parkell) and a second layer of acrylic cement (Lang Dental). Mice were then injected with meloxicam (10 mg/kg) and returned to their homecage for recovery. Mice with GCaMP6s recovered for 6 weeks to allow for viral expression. Mice with array implants recovered for 2 weeks prior to recording.

### In vivo electrophysiology

Adult male mice exposed to ad libitum high fat diet for ten weeks were purchased from Jax and chronically implanted with arrays. Mice were trained on FR operant lever pressing for sucrose prior to in vivo electrophysiological recordings. Mice were recorded during behavior for 16 hours overnight. Physiological signals were recorded with a multi-channel neurophysiology system (Omniplex, Plexon Inc) at 40 kHz and band-pass filtered (150 Hz - 3 kHz) prior to spike sorting. Data analyses were performed with Python and Neuroexplorer (described below).

### Fiber photometry

AAV-DJ-Ef1a-DIO-GCaMP6s was injected and fibers were implanted into the NAc core of D1-Cre or A2A-Cre mice as described above. Six weeks was allowed for recovery and viral expression. A single optic fiber (200 micron core, 0.48 NA, zirconia sleeve connector, Doric Lenses) was used to transmit blue light (475 nm) from a light-emitting diode (Plexbright LED, Plexon) to the brain. The same fiber transmitted emitted light through a dichroic mirror (Doric Lenses). Fluorescence was detected with a photoreceiver (Newport), and signals were amplified and recorded with a digital acquisition system (Omniplex, Plexon).

### Ex vivo electrophysiology

*Slice preparation*. Mice were anesthetized with isoflurance and decapitated. Brains were quickly removed in an ice-cold slurry of sucrose-artificial CSF (aCSF). Sucrose-aCSF comprised the following (in mM): 75 sucrose, 0.5 CaCl_2_, 110 C_5_H_14_CINO, 25 C_6_H_12_O_6_, 25 NaHCO_3_, 7 MgCl_2_, 11.6 C_6_H_8_O_6_, 3.1 C_3_H_3_NaO_3_, 2.5 KCl and 1.25 NaH_2_PO_4_. 220 μm coronal slices were cut using a Leica vibratome (VT 2100), hemisected, and equilibrated at 37 deg C for 45 minutes prior to recording. Slices were then transferred to the recording chamber at 31 deg C and perfused with recirculating aCSF comprising (in mM): 119 NaCl, 2.5 KCl, 1.3 MgCl_2_, 2.5 CaCl_2_, 1.0 Na_2_HPO_4_, 26.2 NaHCO_3_ and 11 glucose and bubbled with carbogen (95% O_2_–5% CO_2_). For intrinsic excitability recordings, the AMPA/NMDA receptor agonist kynurenic acid (2 mM) and the GABA-A receptor antagonist picrotoxin (PTX; 100 uM, Tocris) were present. For mini excitatory post-synaptic currents (mEPSCs), PTX and the voltage-gated sodium channel blocker tetrodotoxin (TTX, 5 uM, Tocris) were present in the bath. *Whole-cell patch clamp recordings*. SPNs were visualized with an upright Olympus X560 microscope and identified by their somal size, and field LED illumination (CoolLED) was used to visualize TdTomato (560 nm). Labeled or unlabelled cells were positively identified as D1^SPNs^ on the basis of TdTomato fluorescence or putatively identified as D2^SPNs^ and targeted for whole-cell recording. Micropipettes were pulled from borosilicate glass capillaries using a pipette puller (Narishige PC-100), and pipette resistance was 6-8 MΩ. To record, electrodes were filled with an intracellular solution with the following (in mM), 130 KGlu, 10 PCr, 4 MgCl_2_, 3.4 Na_2_ATP, 0.1 Na_3_GTP, 1.1 EGTA, 5 HEPES. Osmolarity was adjusted to 289 mOsm, pH 7.3. Recordings were made with a Multiclamp 700B (Molecular Devices) in both voltage and current clamp modes, and analog signals were low-pass filtered at 2 kHz and digitized at 10 kHz with a Digidata 1550 interface (Molecular Devices). Clampex (v 11.4) software was used for data acquisition. In voltage clamp, gigaseal was achieved and cells were held at −70 mV. Reagents were purchased from Sigma unless otherwise specified.

### Data analysis and statistics

#### Closed economy food seeking

Data were written to an SD card on each behavioral device. Custom python scripts were used to quantify effort output and number of trials per session.

#### In vivo electrophysiology

Single and multi-units were identified with principal component analysis (Offline Sorter, Plexon Inc), confirmed and sorted manually, and MANCOVA analyses were used to determine if single units were significantly different from multi-units in the same channel. If p>0.05, spikes were combined as a multi-unit. A Python/Neuroexplorer pipeline was used to generate peri-event histograms of spike activity around reward delivery for each session. Perievent data from all animals was then imported into a single data frame for subsequent analyses in Python. *Normalization*. Signals were Z-scored to a baseline period 10-15 seconds prior to lever press for each perievent histogram. *Classification of units*. Perievent histograms were generated around lever press, and units were either presented collectively (Fig. 2G), or classified via a KMeans clustering algorithm of the three top components contributing to sample variance (Fig. 2I-M). Cluster number was determined using a silhouette analysis. The silhouette score for 3 clusters was 0.42. Principal component analysis and subsequent KMeans clustering was performed on histogram spike profiles scaled to a min-max of 0-1 absolute scale. The top three PCs contributed 54.56 % of the total sample variance. For graphical analysis and quantification, normalized activity was averaged during a baseline period (10 to 15 seconds prior to press), a pre-press period (2 to 6 seconds before press), and during lever press (1 second before and after press is triggered). Baseline, pre-press, and press epochs were chosen in line with previous studies^70^, scaled to encompass a 30 second timeout after a successful trial. ‘Press’ was chosen to encompass neural activity ∼500 ms before and after press. ‘Licking’ epoch was denoted after the release of sucrose and sustained for acerage lick bouts of ∼10 seconds. *Burst analysis*. Bursts were detected with the Burst Analysis tool in Neuroexplorer (0.05 second bin, 3 standard deviations detection threshold), conducted on all units collected during a 60 minute recording in an open arena.

#### Fiber photometry

Photometry signals were corrected for bleaching using a high-pass filter and converted to Z-scores with Python scripts. Perievent histograms were generated around lever press, and averaged during a baseline period (10 to 15 seconds prior to press), a pre-press period (2 to 6 seconds before press), and during lever press (1 second before and after press is triggered). Average normalized fluorescence was compared across periods and between groups.

### Statistics

Comparisons between two samples were conducted with Mann Whitney U tests (Python scipy.stats). Comparisons between more than two samples were conducted with fitted linear models (Python statsmodels). Tukey tests were used for statistical comparison. A p-value < 0.05 was considered statistically significant. For permutation analyses, p-values were calculated by summing the tail of the permutated observations beyond the ground truth data set (percent obese units in each cluster) to determine whether ground truth percentage obese were significantly different from a random distribution of lean and obese classification. Unless otherwise noted, data is presented in scatter plots superimposed on box and whisker plots, with the median and interquartile range represented by the box. All line graphs represent the mean +/- standard error of the mean.

